# EukRibo: a manually curated eukaryotic 18S rDNA reference database to facilitate identification of new diversity

**DOI:** 10.1101/2022.11.03.515105

**Authors:** Cédric Berney, Nicolas Henry, Frédéric Mahé, Daniel J. Richter, Colomban de Vargas

## Abstract

EukRibo is a manually curated, public reference database of small-subunit ribosomal RNA gene (18S rDNA) sequences of eukaryotes, specifically aimed at taxonomic annotation of high-throughput metabarcoding datasets. Unlike other reference databases of ribosomal genes, it is not meant to exhaustively capture all publicly available 18S rDNA sequences from the INSDC repositories, but to represent a subset of highly trustable sequences covering the whole known diversity of eukaryotes. EukRibo strives to include only sequences with verified, up-to-date taxonomic identifications, with a strong focus on protists, and relatively low genetic redundancy, to keep the database compact yet comprehensive. Environmental clone sequences representing previously identified novel diversity are accepted as reference sequences only if they have a precise lineage designation, useful for taxonomic annotation. EukRibo is part of a suite of public resources generated by the UniEuk project, which all follow a common taxonomic framework for maximal interoperability. The high level of taxonomic accuracy of EukRibo allows higher confidence in the taxonomic annotation of environmental metabarcodes, and should facilitate identification of new eukaryotic diversity at various taxonomic levels. The database is currently in version 2, and all versions are permanently stored and made available via the FAIR open platform Zenodo. It is our hope that EukRibo will help ongoing curation efforts of other 18S rDNA reference databases, and we welcome suggestions of corrections and new features to be included in subsequent versions.

## Introduction

High-throughput sequencing of amplicons of genetic diversity markers generated from environmental DNA extractions has become a routinely used approach (typically referred to as “metabarcoding”) to explore the diversity of eukaryotes in all of the planet’s ecosystems (Amaral-Zettler et al. 2009; Cordier et al. 2022; de Vargas et al. 2015; Mahé et al. 2017; Massana et al. 2015). Through reliance on a few relatively well controlled markers such as the key hyper-variable regions V4 and V9 of the small-subunit ribosomal RNA gene (18S rDNA), it is possible to access a much wider diversity of organisms than that which can be isolated and studied in culture conditions. Such environmental surveys have led to a striking improvement in our understanding of the microbial component of the eukaryotic diversity in all habitats. They extend our knowledge of the diversity and geographic distribution of known taxa (Berney et al. 2013; Chambouvet et al. 2014; Egge et al. 2015; Ward et al. 2018), but also reveal the existence of yet unknown lineages (Bass & Cavalier-Smith 2004; Bass et al. 2009; Massana et al. 2014).

Biological interpretation of metabarcoding datasets critically depends on reference genetic databases that allow matching operational taxonomic units / amplicon sequence variants (OTUs/ASVs) with the organism from which they were most likely obtained, by comparison with known gene sequences from identified taxa. This taxonomic identification is what makes it possible to make inferences about the biology and ecological preferences of the organisms that were present in the sampled habitats without actually observing them directly. The two main existing reference databases of 18S rDNA sequences are PR2 (Guillou et al. 2013) and SILVA (Quast et al. 2013). Apart from their use for taxonomic annotation, one of their main goals is also to exhaustively capture all 18S rDNA sequences available in the INSDC repositories. That includes thousands of Sanger-sequenced clones from environmental surveys of eukaryotic diversity. It is important to note that such clone sequences themselves require taxonomic annotation against “primary” reference sequences from described organisms or by phylogenetic inference before they can be used as “secondary” reference sequences to help annotate metabarcoding datasets, and that represents a major taxonomic curation effort.

For the taxonomic annotation of metabarcoding datasets, inclusion of imprecisely or incorrectly annotated environmental clone sequences in reference databases represents a risk of misrepresenting the diversity present in the metabarcoding dataset. There will be a proportion of ASVs in the dataset that have as their closest match in the reference database one of these imprecisely or incorrectly annotated environmental clone sequences. If the annotation was incorrect, these ASVs will be attributed to the wrong taxon. If the annotation was merely imprecise, the ASVs will not be annotated as precisely as they could have been. For instance, they may be annotated as dinoflagellates, but without any further information. But if the imprecisely annotated clone had been excluded from the reference database, it is quite possible that the next closest match would have been a reference sequence identified to the species level, therefore providing a much better taxonomic annotation. Estimation of the phylogenetic novelty in metabarcoding datasets is also highly influenced by the quality of the reference database. ASVs closely matching environmental clones will be considered as “known” because they are represented in the reference database, but in reality they remain of unknown phenotype and with sometimes very limited ecological information; they should best be considered unknown for all intents and purposes. Reference databases including high numbers of environmental clones used as “secondary” reference sequences therefore favour an underestimation of the proportion of ASVs of unknown phenotype even when properly annotated at higher taxonomic levels.

UniEuk is an open, inclusive, community-based and expert-driven international initiative to build a consensus-based, adaptive universal taxonomic framework for eukaryotes (Berney et al. 2017). A key aspect of the UniEuk endeavour is to achieve a taxonomic system that is strictly phylogenetic - using as a starting point the most recent eukaryotic classification (Adl et al. 2019) - and therefore directly applicable to annotation of genetic data. UniEuk comprises three complementary modules, EukMap, EukBank, and EukRef, which use phylogenetic markers, environmental metabarcoding surveys, and expert knowledge to inform the taxonomic framework. EukMap is the core UniEuk module (available at https://eukmap.unieuk.org/), an online resource allowing navigation of the growing UniEuk taxonomy and in its next development phase a direct edition of the taxonomy by members of the research community under supervision by relevant taxonomy experts. EukBank is a module designed to aggregate existing metabarcoding datasets of a given genetic marker into a centralised and publicly available metadataset using a portable analysis pipeline (Berney et al., manuscript in prep.). EukRef is a module designed for taxonomic curation of the 18S rDNA sequences available in the INSDC databases, in particular Sanger-sequenced environmental clones (del Campo et al. 2018).

The EukRef activity is most relevant to the optimisation of the taxonomic annotation of reference sequences in the PR2 and SILVA databases. However this is a very time-consuming process depending largely on hands-on workshops during which sequence curation is achieved on a lineage by lineage basis. Meanwhile, the EukBank module moved forward with the compilation of a first metadataset of metabarcoding datasets targeting the V4 region of the 18S rDNA. One of the main goals of EukBank is to help highlight potential new eukaryotic diversity at various taxonomic levels, both outside and within known groups, that could then be incorporated in EukMap. For this, a reference database compatible with the starting UniEuk taxonomy and of high taxonomic accuracy was required. The timeline of completion of the EukRef curation was incompatible with the immediate needs of EukBank, so that we couldn’t directly use PR2. Instead we decided to build EukRibo, a reduced, PR2-derived, high-quality alternative reference database that would benefit from all the work already done as part of building the UniEuk taxonomy.

### The EukRibo reference database

EukRibo is a manually curated, public reference database of 18S rDNA sequences of eukaryotes. It is provided as one of the public resources accompanying EukBank and it is also closely linked to another UniEuk-related resource - EukProt, a database of genome-scale predicted proteins across the diversity of eukaryotes (Richter et al. 2022). EukRibo does not have the vocation to ever be exhaustive in terms of inclusion of available 18S rDNA sequences in the INSDC repositories. Its main goal is rather to represent a core set of taxonomically well-curated 18S rDNA sequences comprehensively representing all known eukaryotic diversity, that can be used for the taxonomic annotation of high-throughput metabarcoding datasets. It represents a critical stepping stone in the overall UniEuk endeavour that was used to annotate the first version of EukBank, and helped improve the starting UniEuk taxonomy based on 18S rDNA phylogenetic information, and start to link genetic data to taxa and their metadata in EukMap.

To meet its requirements in relation to these various UniEuk activities, the EukRibo reference database was built with the four main following characteristics in mind: (1) a comprehensive phylogenetic coverage; (2) UniEuk-compatible taxonomy strings; (3) a high taxonomic accuracy; and (4) limited sequence redundancy. We expand upon each of these characteristics and their implications below. In addition to its links with other UniEuk resources, it is also our hope that EukRibo will facilitate ongoing curation efforts of the taxonomic annotations in the PR2 and SILVA reference databases. EukRibo remains a work in progress; we welcome suggestions of corrections and new features to be included in subsequent versions.

#### A comprehensive phylogenetic coverage

EukRibo strives to include representative sequences covering all the known eukaryotic diversity (keeping up-to-date with the most recently described taxa and newly characterised deep-branching lineages). With this main goal in mind, the generation of EukRibo was achieved in 6 major steps; the full procedure is described in detail in the Methods section below. Briefly, we used as a starting point PR2 version 4.12.0 (released in August 2019, and containing 183,944 entries / 170,675 unique INSDC sequences) and selected a subset of 44,132 PR2 sequences for which we could ascertain the taxonomic annotation and used it as a starting point. To these we added 2,213 INSDC sequences absent in PR2 version 4.12.0 (mostly because they were recently submitted and not yet included in that version of PR2), and provided UniEuk-compatible taxonomy strings for all entries. This led to a final set of 46,345 reference sequences, including 1,523 environmental clones with a defined lineage identification.

This version 1 of EukRibo, with taxonomy strings that were fixed as of October 2020, was used for the taxonomic annotation of the first EukBank metadataset based on the V4 region of the 18S rDNA. The database is currently in version 2; the main update for this version was to make it taxonomically compatible with version 3 of the EukProt database (https://doi.org/10.6084/m9.figshare.12417881.v3), with taxonomic revisions as of July 2022 (see Availability and versioning below for more details on both versions). Apart from taxonomic updates, version 2 contains the exact same selection of sequences as in version 1, except for the addition of one sequence from the genus *Meteora* (Galindo et al. 2022; Hausmann et al. 2002), the last remaining known eukaryotic lineage for which an 18S rDNA was previously not available (Fig. 1). At this taxonomic level *Meteora* is the only lineage that has been sequenced after EukRibo version 1 was fixed and that is why we decided to add it to version 2. However within all the groups already present in EukRibo some important new diversity has been characterised since, and adding the relevant sequences to EukRibo will be our major task for the future version 3.

**Figure 1.**
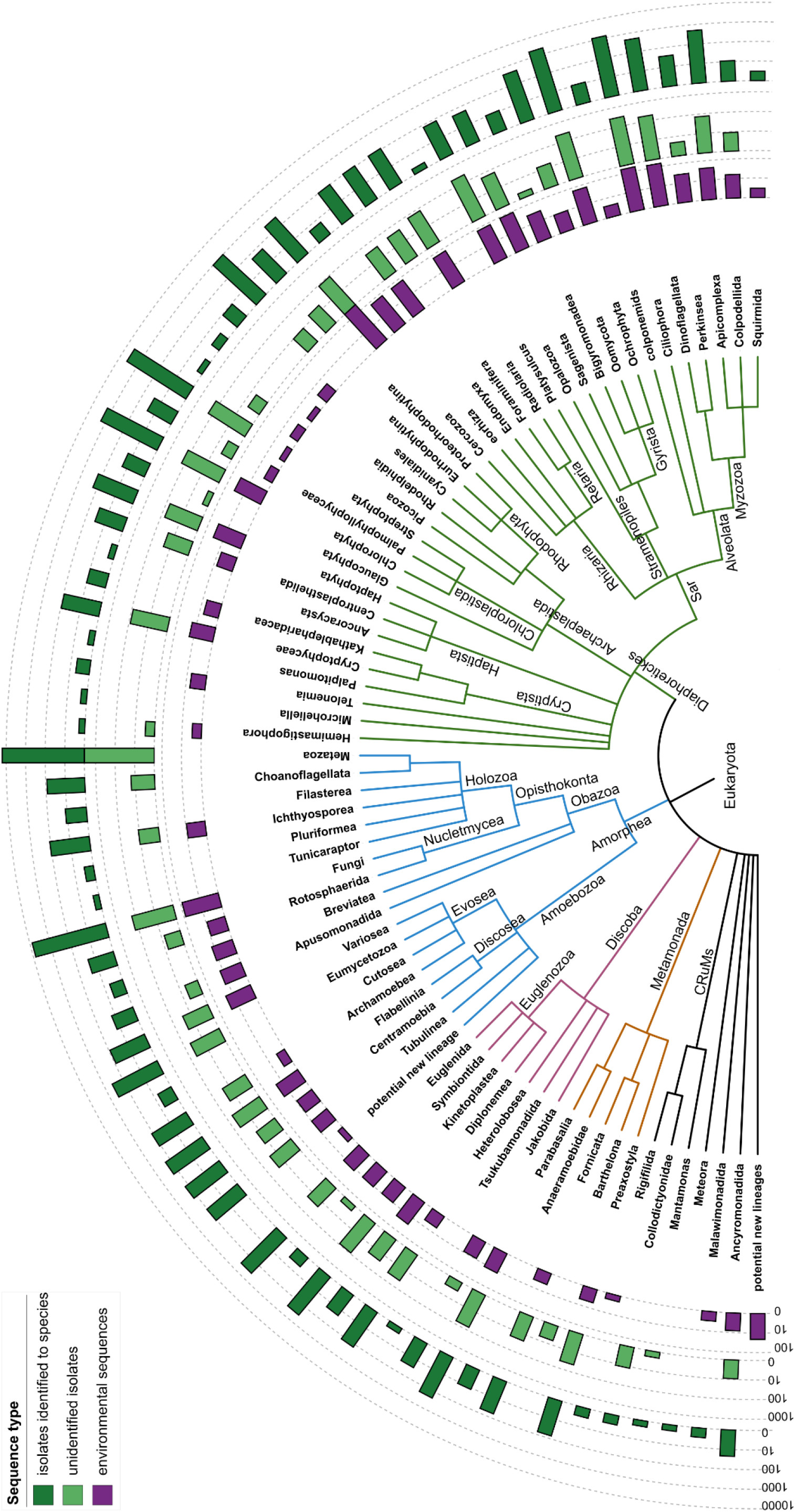
Schematic tree of eukaryotic diversity based on the UniEuk taxonomy strings down to the taxogroup1 level, showing the distribution of the 46,346 reference sequences included in EukRibo version 2 (38,514 from isolates identified to species, 6,304 from unidentified isolates, and 1,528 environmental clones). The position of the eukaryotic root is unresolved, as indicated by a basal polytomy. The figure was created with iTOL (Letunic & Bork 2019).

#### UniEuk-compatible taxonomy strings

All reference sequences in EukRibo were given a UniEuk-compatible taxonomy string to maximise interoperability of the database with UniEuk modules (in particular EukBank and EukMap) and other UniEuk-related public resources such as the EukProt database (Richter et al. 2022). A major difference between this taxonomic system and the one typically used in resources like PR2 is that the taxonomy strings are not based on a fixed number of ranks (typically 8 or 9), but on a free, unlimited number of taxonomic levels, in order to match phylogenetic evidence as closely as possible. In the case of reference databases, this provides end-users more information and flexibility, and when used for taxonomic annotation of genetic datasets, will lead to more precise results. However we recognise that this system can also make it more difficult to summarise results of downstream analyses. Non-specialists cannot guess from taxonomy strings with unlimited ranks which ones are comparable phylogenetically or ecologically. This issue has always been a major rationale for fixing the number of ranks in a resource like PR2. To circumvent this limitation, all EukRibo reference sequences were binned into strictly monophyletic clades at three deep taxonomic levels (also used in EukProt) that we arbitrarily named “supergroup”, “taxogroup1”, and “taxogroup2”. We hope that these three metadata fields will help end-users whenever it is useful to distribute eukaryotic diversity into taxonomic categories of roughly equivalent phylogenetic depth or ecological relevance.

In the context of UniEuk resources, the groupings called supergroups comprise all deep-branching eukaryotic lineages of a phylogenetic depth equivalent to some of the first proposed, “classic” supergroups such as Opisthokonta, Alveolata, Rhizaria, and Stramenopiles. In other words, UniEuk supergroups roughly correspond to the deepest achievable resolution of eukaryotic relationships with 18S rDNA phylogenies, and as a result are expected to be extremely stable through time. They do not necessarily match the highest-level groupings of eukaryotes currently recognized based on phylogenomic evidence, such as Amorphea and Diaphoretickes (Adl et al. 2019) - although these groupings are, of course, present in the complete taxonomy strings. There are currently 36 recognised eukaryotic lineages that were attributed a supergroup level, all of which are represented in EukRibo (Fig. 1). In addition EukRibo includes 9 environmental lineages representing new diversity at a potential supergroup level. UniEuk supergroups are highly variable in relative diversity, ranging from lineages consisting of a single, orphan genus (e.g. *Mantamonas, Meteora, Ancoracysta, Tsukubamonas*), to an assemblage as gigantic as Opisthokonta (containing the multicellular Metazoa and Fungi).

The eight largest supergroups (Opisthokonta, Amoebozoa, Chloroplastida, Rhodophyta, Alveolata, Rhizaria, Stramenopiles, and Euglenozoa) are further subdivided into 3 to 8 taxogroup1 subclades. The larger of these are further subdivided into usually more numerous taxogroup2 subclades, while supergroups of intermediate diversity are directly subdivided into such taxogroup2 subclades. Within a supergroup, the taxogroup1 and taxogroup2 levels should correspond to lineages of relatively equivalent evolutionary or ecological relevance based on current knowledge, although they were more arbitrarily defined than supergroups. As an example, diatoms form taxogroup2 Diatomeae, which is part of the taxogroup1 Ochrophyta within the supergroup Stramenopiles, while flatworms form taxogroup2 Platyhelminthes, which is part of the taxogroup1 Metazoa within the supergroup Opisthokonta. Smaller, ecologically and morphologically more homogeneous supergroups (such as Apusomonadida or Hemimastigophora) or taxogroup1 clades (such as Cyanidiales in red algae and Colponemidae in Aloveolata) are not subdivided further; in such cases the taxogroup2 level is the same as the supergroup or taxogroup1 level.

#### A high taxonomic accuracy

Another main goal of EukRibo was to focus on quality rather than exhaustivity. As described in the Methods below, two main types of verification were performed to ensure the best possible accuracy of our taxonomic curation. First, in many lineages we ran ad-hoc phylogenetic analyses based on alignments of candidate reference sequences to check that their taxonomic annotation matched their phylogenetic position, relying in parallel on our knowledge of INSDC records and NCBI taxonomy issues and any relevant taxonomic literature known to us. Second, we performed a self-annotation of EukRibo that made us very confident about our taxonomic annotations down to the taxogroup2 level. At lower levels the curation quality is currently significantly higher for protists, because they are the focus of the UniEuk project as a whole. In groups outside of the current scope of UniEuk (Metazoa, Fungi, and Embryophyta) taxonomy strings were not checked in detail beyond making sure sequences were attributed to the right lineages just below taxogroup2 level. Within protists, manual taxonomic curation between the taxogroup2 and genus levels was done extensively in all smaller lineages with fewer reference sequences where high taxonomic accuracy was particularly key and easier to ascertain phylogenetically. However in particularly diverse groups like Diatomeae and Chlorophyceae, the level of phylogenetic curation needed to validate all intermediate taxonomic levels (e.g. orders and families) can only be achieved with the EukRef pipeline. This is work in progress; in the meantime rather than risking inclusion of incorrect groupings in our taxonomy strings we deliberately decided to skip some intermediate taxonomic levels in such groups, so that their taxonomy strings can jump directly from class or order to genus.

Our focus on taxonomic accuracy was also reflected in the very strict selection process that we used to validate the inclusion of environmental clones as reference sequences in the database. We mostly selected clone sequences representative of undescribed lineages that had already been identified in previous diversity studies, and ideally attributed a precise environmental lineage identification, useful for taxonomic annotation - such as the various MAST lineages in the Stramenopiles (Massana et al. 2014) for which some ecological information is at least available even if they remain of unknown phenotype. A few previously unrecognised environmental lineages confidently identified in our phylogenetic analyses were also included and given informal names. Much more rarely orphan clone sequences were also considered if they added useful genetic diversity within a phylogenetically well-defined and phenotypically characterised protist lineage.

#### Limited sequence redundancy

Our goal with EukRibo was not just to focus on quality rather than exhaustivity, but actually to strive to limit sequence redundancy as much as possible to achieve a more compact (yet still phylogenetically comprehensive) reference database. So far elimination of this redundancy has mostly been done manually at the species level (especially in animals, land plants, and higher fungi) and it remains imperfect. Still, of the 116,471 candidate reference sequences that were removed at step 3 of the construction of EukRibo (see the Methods below), 51,152 were from isolates, the vast majority of which corresponded to organismal redundancy at the species level. In subsequent versions of EukRibo we plan to tackle the remaining sequence redundancy in a more systematic way using clustering of the reference sequences, especially in the multicellular clades mentioned above. The rationale behind this reduction of sequence redundancy is to optimise EukRibo for the kind of downstream applications that are actually impeded by the sequence exhaustivity of the PR2 and SILVA databases. In particular, we would like EukRibo to ultimately comprise the smallest possible set of reference sequences necessary and sufficient to perform highly accurate taxonomic annotation of metabarcoding datasets at higher taxonomic levels, and allow identification of new phylogenetic diversity at various taxonomic levels. In the following section we discuss in more detail the usefulness of smaller yet comprehensive and taxonomically accurate reference databases in the context of the taxonomic annotation of high-throughput environmental datasets.

### Using EukRibo for taxonomic annotation of metabarcoding datasets

Classically the taxonomic annotation of ASVs from metabarcoding datasets against a reference database has been based on the taxonomy of the reference sequence(s) with the highest similarity to the ASV, as estimated by BLAST algorithms (Altschul et al. 1997) or with tools like VSEARCH (Rognes et al. 2016). A minimum similarity threshold is generally also applied (typically around 80-85%), under which the ASV is considered taxonomically unassigned. This approach gives excellent results for ASVs that exactly or very closely match reference sequences (>99% similarity). And below a certain threshold, it is correct to consider the ASV unassigned. But the main limitation of this approach is that in between these two extremes usually lies a huge novel genetic diversity that gets incorrectly assigned a full taxonomy string down to species level, even though it clearly corresponds to different organisms absent from the reference database. An ASV with e.g. 93% maximum similarity to a reference sequence cannot possibly be the same species, but most likely shouldn’t be considered totally unassigned either. But where should the taxonomic annotation stop - at genus level, family level, order level?

Within the UniEuk effort, one of the main goals of the EukBank resource is to help highlight potential new eukaryotic diversity at various taxonomic levels, both outside and within known groups. This is why the V4 region of the 18S rDNA was the diversity marker chosen for version 1 of EukBank: in eukaryotes the V4 region is around 375-380bp. long on average, and although it contains several hyper-variable secondary structure elements (Wuyts et al. 2000) that allow taxonomic distinctions down to genus level (or in some lineages even down to species level), it also contains more conserved regions with a good phylogenetic signal at higher taxonomic levels. Therefore the V4 region should be an ideal marker to achieve taxonomic annotations and detect novel diversity in a more phylogenetically-informed way than with genetic markers that may be better metabarcodes at species level but lack phylogenetic signal at higher levels, such as the ITS regions used in Fungi (Schoch et al. 2012).

To test this hypothesis we decided to use the set of reference sequences included in EukRibo to check whether sequence comparisons of the V4 region allowed detection of “natural” similarity gaps between phylogenetic groups, correlated with a V4 phylogenetic signal. For this we used the supergroup, taxogroup1, and taxogroup2 bins that we attributed to all sequences. As part of the creation of EukRibo (see Methods below) we performed a pairwise alignment of each reference sequence against all others. We used the resulting similarity scores and for each reference sequence, we plotted its best similarity score to another sequence in the same supergroup against its best similarity score to any sequence in a different supergroup, then repeated the same process for the taxogroup1 and taxogroup2 levels (Fig. 2). The resulting plots show that “natural” similarity gaps do exist in the V4 region between phylogenetic groups. At supergroup level for instance, a sequence that has a similarity score >99% to at least one other sequence in the same supergroup will never have a maximum similarity above 90% to any sequence in a different supergroup. Unsurprisingly the similarity gap decreases at lower taxonomic levels. But even at taxogroup2 level it remains visible, with a minimum similarity gap between 1.5 and 2% even for the closest taxogroup2 bins - e.g. some protostome animals, some streptophyte algae, or some haptophyte lineages.

**Figure 2.**
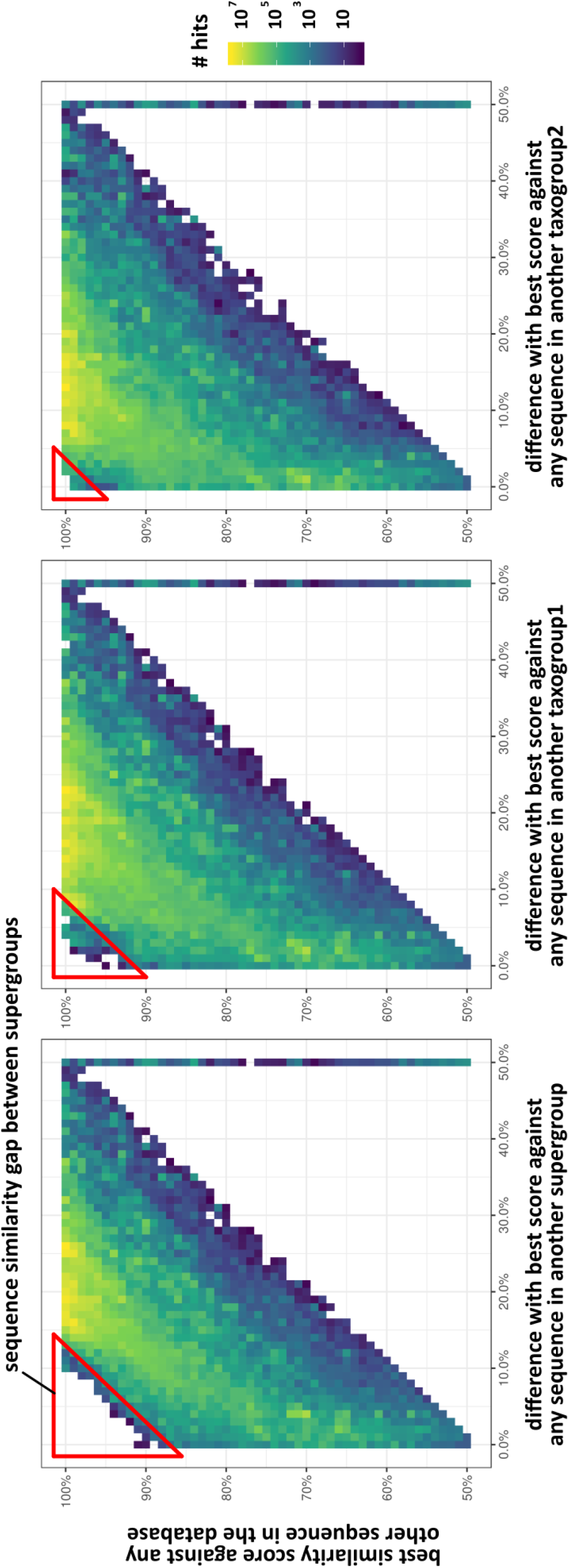
Plots of the similarity scores resulting from the self-annotation of EukRibo, illustrating the existence of maximum similarity gaps between the supergroup, taxogroup1 and taxogroup2 taxonomic levels that can be applied to taxonomic annotation of ASVs from metabarcoding datasets. The extracted V4 region of each EukRibo sequence was compared to all others by pairwise alignment using VSEARCH. The similarity score of each pairwise comparison was mapped along the Y-axis according to its best similarity score against any other sequence in the database (i.e. including those from the same lineage), while the X-axis shows the % difference with the best similarity score against any sequence in another supergroup, taxogroup1, or taxogroup2, respectively.

In light of this result we conceived a very straightforward, fast yet efficient taxonomic annotation approach that allows confident assignment of ASVs to any number of pre-defined, hierarchically organised taxonomic bins of reference sequences, with the only condition that there should be an empirically demonstrable similarity gap between them. The approach is simply based on the VSEARCH similarity scores of a pairwise alignment of each ASV against all reference sequences. It uses the observed similarity gaps between the best hits in different taxonomy bins to determine whether an annotation to that taxonomy bin is validated (because the similarity gap is equal to or higher than that empirically observed between reference sequences alone) or if it is rejected. The approach and the specific similarity gaps we recommend to use for supergroup, taxogroup1 and taxogroup2 annotations for the V4 region based on EukRibo reference sequence comparisons are described in detail in the Methods section below.

This approach is made possible by (1) the high taxonomic quality of the EukRibo reference database - ensuring the existence of empirically demonstrable similarity gaps between taxonomic bins; and (2) its small size - making the approach scalable even if the metabarcoding dataset(s) to annotate increase in size. Indeed with large metabarcoding datasets pairwise alignment of each ASV against all reference sequences would quickly get too computationally intense if the reference database was too large. We tested our approach on the EukBank ASVs and got promising results (manuscript in prep.); the idea was to ensure the best possible compromise between achieving high confidence in the attribution of ASVs to higher taxonomic levels, and maintaining a high speed of analysis given that a metadataset such as EukBank is expected to continue growing in the future.

Having UniEuk taxonomic pathways with an unlimited number of ranks also makes this approach highly flexible. Because we were mainly focused on confidently assigning ASVs to higher-level lineages, we currently chose to stick to three levels supergroup, taxogroup1 and taxogroup2, for which we empirically verified the existence of sequence similarity gaps. This suggests that these levels would also be well-suited for probabilistic taxonomic annotation approaches relying on principles from machine learning to reduce over classification errors, such as IDTAXA (Murali et al. 2018). But the approach could be applied to any combinations of taxonomic bins at any taxonomic level as long as appropriate similarity gaps have been empirically determined using comparisons of the reference database against itself. Another goal for subsequent versions of EukRibo will therefore be to automatise the identification of the taxonomic levels within the full taxonomy strings that meet this requirement. This will make it possible to highlight the specific subset of taxonomic levels in any taxonomy string with unlimited ranks that is most relevant for taxonomic annotation purposes, either with our newly conceived approach or with classifiers like IDTAXA.

For these reasons we believe EukRibo to be the ideal resource for a rapid yet efficient taxonomic annotation of large metabarcoding datasets at higher taxonomic levels and to detect ASVs representing new genetic diversity. Importantly however, EukRibo is not meant to ever replace the PR2 or SILVA reference databases. EukRibo is deliberately meant to only ever include a phylogenetically comprehensive but small subset of all 18S rDNA sequences present in INSDC. By contrast, thanks to their exhaustive inclusion of 18S rDNA sequences (in particular environmental clones), PR2 and SILVA remain the best resources to assess when and where a certain lineage or ASV has been observed before, providing key insight into the geographic distribution and ecological preferences of the organisms behind the sequences. Ultimately we could imagine EukRibo being absorbed by SILVA or PR2 in the form of additional metadata fields highlighting the smallest subset of reference sequences necessary and sufficient for taxonomic annotation and detection of new diversity. In the meantime we would encourage parallel use of these public resources to maximise biological inferences from metabarcoding datasets.

## Methods

### Construction of the EukRibo database

The generation of EukRibo version 1 was achieved in 6 major steps detailed below, using as a starting point PR2 version 4.12.0 (released in August 2019, and containing 183,944 entries / 170,675 unique GenBank sequences).

*Step 1 - preselection of PR2 sequences to be considered* Automatic elimination of entries that (1) do not correspond to nuclear eukaryotic 18S rDNA sequences, using the following PR2 fields: “kingdom”, “gene”, “organelle”, and “gb_locus”; (2) were not direct submissions to NCBI, EMBL-EBI/ENA or DDBJ, using the meaning of INSDC accession number prefixes; and (3) exist in duplicates in PR2 (e.g. with and without introns), keeping only unique accession numbers. Following this step 22,472 entries were removed, with 161,472 entries / sequences going through to the next step.
*Step 2 - preliminary taxonomic curation based on PR2 information*
  A. Extraction of the unique PR2 taxonomy strings present in these 161,472 sequences and translation into preliminary UniEuk taxonomy strings using a manually constructed conversion table.
  B. All sequences flagged according to their origin (94,724 entries from isolates that would *a priori* be kept, and 66,748 entries that are environmental clones and would *a priori* be excluded).
  C. Taxonomic binning of the sequences into ca. 50 clusters of “supergroup” level (eukaryotic lineages of a phylogenetic depth corresponding to e.g. Opisthokonta, Chloroplastida, or Alveolata) and ca. 360 clusters of “taxogroup2” level (eukaryotic lineages of a phylogenetic depth corresponding to e.g. Arthropoda, Mamiellophyceae, or Colpodea), sometimes with an intermediate “taxogroup1” level (eukaryotic lineages of a phylogenetic depth corresponding to e.g. Metazoa, Chlorophyta, or Ciliophora); from then onwards work was done at a supergroup, taxogroup1, or taxogroup2 level.
*Step 3 - in depth manual taxonomic curation*
  A. Manual verification and edition of the preliminary UniEuk taxonomy strings automatically inferred from PR2, based on (1) our knowledge of INSDC records and NCBI taxonomy issues, (2) all relevant literature known to us, and (3) ad-hoc phylogenetic analyses. For Metazoa, Fungi and Embryophyta, the PR2 taxonomy was globally trusted. Intermediate phylogenetic levels absent in PR2 were automatically extracted from the full NCBI taxonomy strings (which also allowed a verification of the PR2 annotations); only levels that were deemed phylogenetically informative enough were kept.
  B. Manual elimination of some of the organismal redundancy at species level, especially in animals, land plants, and higher fungi.
  C. Manual addition of (mostly recently submitted) INSDC sequences missing in PR2 version 4.12.0 but representing important taxonomic diversity; this included five supergroup-level clades such as Hemimastigophora.
  D. Manual selection of good quality environmental clone sequences that are representatives of previously identified (environmental) diversity and that could be attributed an informative taxonomy string after validation by phylogeny.
  E. Manual addition of 40 representative nucleomorph sequences from the lineages Cryptomonadales and Chlorarachniophyceae. Nucleomorph sequences typically do get amplified alongside their nuclear counterparts in eukaryotic environmental surveys; it is necessary to include representative nucleomorph sequences in EukRibo to annotate the corresponding ASVs, which can then if needed be eliminated before further analyses. Following this step 116,471 sequences were removed (51,152 from isolates and 65,319 environmental), 45,001 sequences were kept (43,572 from isolates and 1429 environmental), and 2,228 missing sequences were added (2,138 from isolates and 90 environmental), with 47,229 sequences going through to the next step.
*Step 4 - extraction of the V4 region from all sequences* Because the primary goal of EukRibo is to be used to annotate the EukBank metadataset of available V4 metabarcoding datasets, we decided that all sequences included in EukRibo should contain the V4 region. This step was done automatically whenever both primer sequences were found in the sequences (allowing up to 20% mismatches), with visual inspection to check that the V4 region was correctly extracted. Sequences where one or both of the primer sequences could not be found automatically were checked manually. Sequences that do not include the V4 region were eliminated (with replacement by other taxonomically equivalent sequences whenever possible). Because exhaustive taxonomic coverage is a crucial feature for EukRibo, sequences with a slightly incomplete V4 region were kept if they were phylogenetically useful - i.e. if they were the only available representatives of a certain taxonomic lineage. We allowed up to 50 missing positions in the relatively conserved area at the 5’ end of the V4 region (for an average fragment length of about 375-380 bp.); no sequence incomplete at the 3’ end of the V4 region was included. Following this step 967 sequences were removed and 84 sequences were added, with 46,346 sequences going through to the next step.
*Step 5 - checking the internal congruence of EukRibo* A verification of the validity of the supergroup, taxogroup1, and taxogroup2 binning and of the general congruence of the full taxonomy strings was achieved by taxonomic annotation of the database against itself, using pairwise alignment of each reference sequence against all others. Taxonomic conflicts (resulting from contaminations or from misidentifications that had not been detected during Step 3, mostly in non-protist groups) were eliminated. A few useful reference sequences newly identified after Step 3 were also added at this stage. The self-annotation was then repeated to make sure that no new conflict could be detected. Following this step 42 sequences were removed and 41 sequences were added, leading to the final total of 46,345 V4-containing sequences included in EukRibo version 1.
*Step 6 - extraction of the V9 region from all sequences* This step was performed to highlight the subset of EukRibo sequences that contain both the V4 and V9 regions. This will make it possible for users to compare with the exact same set of reference sequences the respective coverage of eukaryotic diversity achieved with V4 and V9 when both markers are targeted in the same samples. This step was done as described above for the V4 region. Again, sequences with slightly incomplete V9 regions were kept to maximise taxonomic coverage. In this case we allowed up to 30 missing positions in the relatively conserved area at the 3’ end of the V9 region (for an average fragment length of about 130-135 bp.); no sequence incomplete at the 5’ end of the V9 region was included. A self-annotation step was also performed as described above to verify that the taxonomy inferred from the V9 region was compatible with the full taxonomy strings. It allowed identification of a few chimeric sequences (where the V9 region does not originate from the same organism as the V4 region); for these the V9 region was not kept. Because many 18S rDNA sequences in the INSDC repositories stop before the 3’ end of the gene, only about 75% of EukRibo sequences contain a sufficiently complete V9 region, leading to the final total of 34,438 V9-containing sequences included in EukRibo version 1.

### A new simple approach to annotate metabarcoding datasets

Taking advantage of the small size and high taxonomic quality of the EukRibo reference database, we conceived a very straightforward new taxonomic annotation approach that makes it possible to confidently assign ASVs to any number of pre-defined, hierarchically organised taxonomic bins of reference sequences - in our case the supergroup, taxogroup1 and taxogroup2 levels that we attached to all EukRibo reference sequences.

*Step 1 - classic annotation of ASVs against EukRibo* All ASVs are annotated using a classic approach by pairwise alignment against each and every sequence in EukRibo version 1 using the stampa pipeline (https://github.com/frederic-mahe/stampa/), which relies on VSEARCH similarity scores. Pairwise-similarity percentages are calculated for all sequence comparisons, analysed, and stored for the next step. Taxonomic assignment is based on the taxonomy of the last common ancestor of the co-best hits among all reference sequences.
*Step 2 - attributing ASVs to the supergroup, taxogroup1 and taxogroup2 levels* Final assignment of ASVs to the supergroup, taxogroup1 (if relevant) and taxogroup2 levels is done as follows:

- ASVs with at least 99% similarity to a reference sequence are considered correctly assigned at all 3 levels. This is based on the observation that the similarity gap between any pair of EukRibo sequences from different taxogroup2 levels is always greater than 1.5% (Fig. 2).
- For similarity values below 99%, assignment to a supergroup is validated if no assignment to an alternative supergroup is observed within a 9% similarity gap margin. This is based on the observation that the minimum similarity gap between any pair of EukRibo sequences from different supergroup levels is always at least 9% (Fig. 2). ASVs not meeting this criterion are considered unassigned at the supergroup level (and consequently at the two taxogroup levels as well).
- Once assigned to a supergroup, assignment to a taxogroup1 (if relevant) is validated if no assignment to an alternative taxogroup1 is observed within a 6% similarity gap margin. Again this is based on the observation that within a supergroup the minimum similarity gap between any pair of EukRibo sequences from different taxogroup1 levels is around 5%. ASVs not meeting this criterion are considered assigned at the supergroup level but unassigned at the taxogroup1 level (and consequently at the taxogroup2 level).
- Assignment to a taxogroup2 is validated if no assignment to an alternative taxogroup2 is observed within a 3% similarity gap margin. Again this is based on the observation that within a supergroup (or taxogroup1, if relevant) the minimum similarity gap between any pair of EukRibo sequences from different taxogroup2 levels is between 1.5 and 2%. ASVs not meeting this criterion are considered assigned at the supergroup level (and taxogroup1, if relevant) but unassigned at the taxogroup2 level.
- Conservatively, ASVs with an overall best similarity score to any reference sequence lower than 70% are considered as unassigned at all levels irrespective of the similarity gap between the best and second best supergroup assignments. This threshold was chosen to make sure that it remains above mere random similarity.

Note: the lower the maximum similarity score to a reference sequence is, the less reliable these assignments will become. We’d advise caution in accepting assignments without any phylogenetic verification when the maximum similarity to a reference sequence is below 75%. However, we have noticed empirically a very clear difference between ASVs that have close similarity scores (even when above 85%) across many supergroups and/or taxogroups - and likely represent new diversity at that taxonomic level - as opposed to ASVs that have a significantly higher maximum similarity score (even if below 75%) against one supergroup and/or taxogroup than against all others. We interpret this observation as a consequence of the high length heterogeneity across eukaryotes in the V4 region. Pairwise-similarity percentages quickly decrease in presence of taxon-specific insertions (whether they are introns or expansions in hypervariable V4 helices) even between close relatives. But overall, phylogenetically-driven similarity in the conserved parts of the V4 region will still provide a strong enough phylogenetic signal to ensure the existence of significant similarity gaps between groups.

### Availability and versioning

The EukRibo database is permanently stored and made available via the FAIR open platform Zenodo at https://doi.org/10.5281/zenodo.6327890. Each EukRibo release consists of four files: (1) a tsv table containing the taxonomic and other information about the 18S rDNA sequences included in the release; (2) a fasta file containing the full sequences as retrieved from the INSDC repositories; (3) a fasta file containing the variable region V4 extracted from all these sequences (based on the fragment amplified with the Tara-Oceans V4 primers); and (4) a fasta file containing the variable region V9 extracted from the subset of sequences where it is present (based on the fragment amplified with the Tara-Oceans V9 primers).

Version 1 was published in March 2022. This is the starting version of EukRibo that was used for the taxonomic annotation of the EukBank dataset, with taxonomy strings that were fixed as of October 2020. It contains 46,345 reference sequences including a sufficiently complete V4 region; 46,299 with the actual complete V4 region and 46 (about 0.1%) with up to 50 missing positions at the 5’ end of the V4 region. Of these, 34,438 reference sequences also include a sufficiently complete V9 region; 23,226 with the actual complete V9 region and 11,206 (about 33%) with up to 30 missing positions at the 3’ end of the V9 region. The tsv table in version 1 includes the following six fields: (1) “gb_accession” - the INSDC accession number of the sequence; (2)

“UniEuk_taxonomy_string” - the full, UniEuk-compatible taxonomic annotation of the sequence, allowing an unlimited number of levels (going down to strain for isolated organisms or to clone for environmental sequences) as well as informal names for phylogenetically supported clades without formal name; (3) “V9” - indicating the presence (“Y”) or absence (“N”) of a sufficiently complete V9 region in the sequence; (4) the three fields “supergroup”, “taxogroup1”, and “taxogroup2” - corresponding to the binning of the taxa into strictly monophyletic clades of evolutionary significance.

Version 2 was published in July 2022; it contains the exact same selection of sequences as version 1, with the addition of one extra sequence representing the last remaining known supergroup-level eukaryotic lineage for which an 18S rDNA sequence was not previously available, the enigmatic genus *Meteora*. In addition, the taxonomy string and supergroup, taxogroup1 and taxogroup2 binnings of several reference sequences were updated (taxonomic revisions as of July 2022) to make this version of EukRibo taxonomically compatible with version 3 of the EukProt database (https://doi.org/10.1101/2020.06.30.180687). Only 34,432 sequences with a sufficiently complete V9 region are now retained because of 6 previously unrecognised chimeric sequences where the V9 fragment does not originate from the same organism as the V4 fragment. (The new *Meteora* sequence contains the full V4 fragment but does not include a sufficiently complete V9 fragment.)

The tsv table in version 2 now includes twelve fields instead of six. The fields “gb_accession”, “supergroup”, “taxogroup1”, “taxogroup2”, and “UniEuk_taxonomy_string” are the same as in version 1. The following fields are new: (1) “alternative_strain_names” - providing alternative strain/isolate names when known to help cross-linking genetic data coming from the same organism; (2) the two fields “EukProt_ID_same_strain” and “EukProt_ID_different_strain” - providing the ID of EukProt datasets from the same isolate or a different isolate of the same species, respectively; (4) “columns_modified_since_previous_version” - listing pre-existing fields that have a modified content compared to version 1; (5) “remarks” - providing additional information such as presence of an intron in the V9 fragment, or the taxonomic identity of the two parts of chimeric sequences; (6) “V4” - indicating whether the V4 fragment is complete (“yes - complete”) or missing some positions at the 5’ end (“yes - partial”). Finally, the content of field “V9” was amended to include more precise information about whether region V9 is complete (“yes - complete”), missing some positions at the 3’ end (“yes - partial”), or was excluded, with six possible reasons why (“no - missing”, “no - too incomplete”, “no - chimera”, “no - bad quality”, “no - deletion in V9”, “no - Ns in V9”).

## Acknowledgements

CB was supported by grants from the Gordon and Betty Moore Foundation (GBMF5257 & GBMF8908 / UniEuk project) and is grateful to the International Society of Protistologists for additional financial support. DJR was supported by postdoctoral fellowships from the Beatriu de Pinós programme of the Government of Catalonia’s Secretariat for Universities and Research of the Ministry of Economy and Knowledge, and from ”la Caixa” Foundation (ID 100010434), with the fellowship code LCF/BQ/PI19/11690008. This work was supported by the French Government ‘Investissement d’Avenir’ program OCEANOMICS (ANR-11-BTBR-0008) and the ABiMS computing cluster at the Station Biologique de Roscoff, France.

